# Comparison of library preparation and sequencing depths for direct sequencing of *Bordetella pertussis* positive samples

**DOI:** 10.1101/2021.02.14.430694

**Authors:** Winkie Fong, Keenan Pey, Rebecca Rockett, Rosemarie Sadsad, Vitali Sintchenko, Verlaine Timms

## Abstract

Whooping cough, or pertussis, is a highly transmissible respiratory infection caused by *Bordetella pertussis*. Due to the high burden of pertussis, vaccine programmes were introduced internationally and in Australia since the 1950s. This has resulted in a significant decrease of pertussis infections. However, since the 1990s the number of pertussis notifications has increased considerably. Currently circulating *B. pertussis* strains differ in vaccine antigen composition compared to strains that circulated in the pre-vaccination era. These genetic differences are thought to contribute, in part, to the re-emergence of pertussis in Australia and around the world. Whole genome sequencing (WGS) can resolve minute differences in circulating strains and provides unparalleled resolution of vaccine antigens. This high-resolution snapshot can provide clues that enable more targeted public health interventions. However, pertussis is primarily diagnosed with culture-independent diagnostic assays which offer fast turnaround result times and reduced laboratory costs, eliminating the need to culture isolates. Current WGS methods require a cultured isolate, resulting in an absence of *B. pertussis* genome sequences in the post vaccination era. This scarcity has, in turn, limited understanding of currently circulating strains and respective vaccine antigen compositions.

Recent advancements of WGS technologies have allowed direct sequencing of clinical specimens without the need for a cultured isolate. However, recovering reliable sequence data from clinical samples of low bacterial load infections such as *B. pertussis* is a pressing challenge. We sought to increase the yield of *B. pertussis* sequences direct from a clinical sample by evaluating widely available WGS library preparation methods.

We report that the Illumina DNA prep library preparation kit combined with deep sequencing allowed the detection of important surveillance information such as allelic variations in the *B. pertussis* vaccine antigens. Further, our method generates high coverage over the 23S ribosomal RNA of *B. pertussis* enabling macrolide resistance to be easily determined. Overall, this method can improve surveillance of *B. pertussis*, by monitoring changes in vaccine antigens, detecting antimicrobial resistance and guiding Public Health control interventions.

## Introduction

*Bordetella pertussis*, the primary causative agent of whooping cough, is a highly contagious respiratory pathogen that causes mild symptoms in adults but severe infection with a high mortality rate in infants. Global vaccination efforts have significantly reduced pertussis mortality, however, pertussis remains endemic globally, even in countries with high vaccine coverage^1^. The cause of the recent resurgence is hypothesised to include waning host immunity^2^ and vaccine escape^3^. Investigation of the persistence of pertussis has been challenged by the lack of *B. pertussis* isolates. The introduction of highly sensitive and fast PCR-based diagnostics has rendered culture of *B. pertussis* largely redundant, and limited the ability to perform phenotypic strain typing and antibiotic susceptibility testing^4^. The re-emergence of pertussis, along with the emergence of macrolide resistant *B. pertussis*, demonstrates how critical it is that we continue to monitor *B. pertussis* strains, despite a decline in the number of culture isolates.

Whole genome sequencing (WGS) of bacteria with epidemic potential provides highly nuanced data^5, 6^ used in outbreak control where it can identify clusters and potential transmission chains^6^. Read depth, defined as the number of reads over a genome position, and genome coverage, the percentage of read coverage across the genome, are important parameters in direct sequencing protocols as they provide assurance of the accuracy and quality of the consensus sequence. When this high-resolution technology is applied directly to a clinical sample the overwhelming majority of sequences obtained are of human origin. In a respiratory sample, the number of human cells poses the greatest obstacle to resolving sufficient sequencing reads for a low-load pathogen, such as *B. pertussis*^6^. Studies have previously shown that pre-treatment of nasopharyngeal aspirates (NPA) can reduce human DNA levels, and increase target pathogen DNA and therefore pathogen specific sequencing reads^4, 7^. However, despite the increased yield of *B. pertussis* from a saponin treated sample, microbial genome coverage and read depth was still insufficient in providing informative molecular typing data^4^. One way to improve sequence coverage is increase the number of total reads per samples, known as deep sequencing. Deep sequencing aims to increase the read depth and the accuracy of detecting mutations over positions in genes of interest by generating more unique reads across the genome^8^. However, read depth and coverage can also be enhanced by selecting the appropriate library preparation method^9^.

Library preparation kits play a critical role in coverage yield, as the method of fragmentation used can introduce significant biases. For *B. pertussis*, G+C bias is a major limiting factor in transposase and PCR-based library preparation methods as it results in a loss of reads and low coverage over high G+C content areas of the genome^10-13^. Considering *B. pertussis* has an average G+C content of 67%, library preparation methods (e.g., Nextera XT) can bias library construction to human DNA present and not produce libraries as efficiently from *B. pertussis* DNA present in a clinical sample^10-13^. Moreover, in contrast to mechanical fragmentation-based library preparation methods, enzymatic fragmentation contains slightly greater insertion biases due to sequence preferences of the transposase^14-16^. While there is little impact of these insertion biases on rapid, parallel sequencing of isolates, the effect of this bias on sequencing of a high G+C content pathogen directly from clinical samples cannot be ignored and requires further investigation^12, 17^.

This study examines the influence of library preparation on culture-independent sequencing of *B. pertussis*. We compare library preparation kits from Illumina, the most widely used platform in Public Health Microbiology laboratories and assess the read depth and genome coverage of a low-load, high GC pathogen. Further, by comparing libraries at increased sequencing depths we investigate whether accurate genomic information relevant for public health control of *B. pertussis* can be recovered directly from a clinical sample.

## Methods

Nasopharygeal aspirates (NPA) were collected from the Centre of Infectious Diseases and Microbiology – Laboratory Services (CIDMLS), NSW Health Pathology, Westmead Hospital, Sydney. These NPA specimens were submitted for clinical testing of other infectious diseases and were negative for *B. pertussis*. The samples were pooled and 200 μL was taken for baseline extractions with the Qiagen DNeasy Blood and Tissue (QIAGEN, Germany) DNA Extraction Kit (further described below) to confirm the NPA was *B. pertussis* negative by PCR. The pooled NPA were then spiked with 0.5 McFarland suspension of *B. pertussis* (ATCC®9797™ 18323) corresponding to approximately 1.4 × 10^6^ CFU/mL (BP10-2), 1.4 × 10^5^ CFU/mL (BP10-3), and 1.4 × 10^4^ CFU/mL (BP10-4), then stored at −20°C. The specific dilutions were chosen as C_T_ cycles by *IS481* real-time PCR (rtPCR) demonstrate these are within the normal range of clinical samples^18^. Further clinical NPA (n=2) that were positive for *B. pertussis* and collected by CIDMLS with the highest C_T_ cycle were included for standard and deep sequencing. Details of *IS481* PCR are presented in the Supplementary Material.

### Extraction and DNA Quality Control

The spiked and clinical samples were treated with 0.025% Saponin and TurboDNase as previously described^4, 7^. The spiked samples were then extracted with the Qiagen DNeasy Blood and Tissue (QIAGEN, Germany) DNA Extraction Kit for “Purification of Total DNA from Animal Tissues (Spin-Column Protocol)” with modifications. The modifications included lysis of the NPA with Proteinase K at 56°C for 1.5 hours. The DNA extracts were split into four aliquots for use in rtPCR targeting *IS481* and *ERV3* as previously described^4^, and for use in library preparation method comparison (Supplementary Material).

### Library Preparation and Sequencing

The three Illumina (Illumina, USA) library preparation kits selected were; NexteraXT DNA Library Preparation Kit v2.5, TruSeq DNA Nano Library Preparation Kit and the Illumina DNA prep Library Preparation Kit. Libraries were constructed following the manufacturer’s protocol. The two clinical NPA samples were prepared with Illumina DNA prep only. Fragment size and quality was assessed by the Agilent TapeStation 4200 using the High-Sensitivity DNA ScreenTape Assay (Agilent, USA) on TruSeq input DNA pre- and post-sonication, and Nextera XT, Illumina DNA prep and TruSeq post-library construction. Library quantification was assessed using the KAPA (Roche Diagnostics, Switzerland) assay on all libraries for normalisation and final pool concentration. Illumina DNA prep libraries were sequenced on the Illumina NextSeq 500 instrument in conjunction with other routine samples that generate between 3-5 million reads per sample. TruSeq and Illumina DNA prep libraries were sequenced together on the Illumina MiniSeq High-Throughput platform. These were considered as sequenced on a standard level. In addition, Illumina DNA prep libraries were sequenced again on the Illumina NextSeq 500 with a High-Throughput flow cell for deep sequencing to generate around 80 million reads per sample.

### Analysis

#### Quality Control

Analysis began with FastQC^19^(v 0.11.3) to assess the sequencing quality of raw reads, followed by trimming the ends of poor-quality reads with Trimmomatic (v0.36)^20^ using optimised parameters (Leading:3 Trailing:3 SlidingWindow:4:20 Minlen:36).

#### Mapping

Trimmed reads were then mapped to the Human genome GRCh38.p12 (GCA_000001405.27) by Burrows-Wheelers Aligner (BWA v0.7.12) using default settings^21^. Three strains of *B. pertussis* were utilised as mapping reference genomes – Tohama I reference genome (NC_002929.2), ATCC®9797™ 18323 whole genome (NC_018518.1) as this was the strain used to spike the NPA, or B1917 (NZ_CP009751.1) as this is a representative strain of currently circulating *B. pertussis*^22^.

#### Gene and Genome Counts

Percentage of each representative organism (Human and *B. pertussis*) was determined by the number of reads mapped to the respective genomes divided by the number of reads of the total sample.

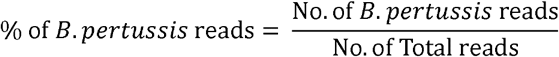

Manual visualisations of coverage across genes of interest such as the 23S ribosomal RNA, vaccine antigen genes: *ptx, prn*, and *fhaB* encoding regions were performed using Qiagen CLC Genomics Workbench 12 (Qiagen, Germany). The list of all genes of interest has been provided in Supplementary Material.

HTSeq (v11.2)^23^ with default ‘union’ settings was used to count the number of unique reads mapped over these genes of interest, mapped reads over multiple genes were discarded. Samtools depth (v1.9)^24^ and BEDtools intersectBed and genomecov (v2.25.0)^25^ were used to calculate the average read depth, gene and genome coverage and plotted with ggplot2 (v3.3.0)^26^.

## Results

The first question this study aimed to address was whether the method of library preparation had any impact on the sequencing efficiency of a high G+C, low load microbial pathogen such as *B. pertussis*.

### NexteraXT DNA Library Preparation using standard sequencing

As expected, the percentage of *B. pertussis* reads identified decreased as CFU decreased (Table 1). Genome mapping to both *B. pertussis* (Tohama I and ATCC®9797™ 18323) genomes show only 56.5 ± 0.5% of the genomes are covered by these reads. Deeper analysis into coverage over the vaccine antigen and 23S ribosomal RNA showed poor coverage across the vaccine antigen genes, with BP10-2-1 failing (zero reads) to present reads to 10 out of 14 genes and BP10-2-2 failing 3 out of 14. BP10-3-1 contained reads to all genes, 170 reads mapped to *fhaB* which accounted for 85.6% of the gene. BP10-3-2, BP10-4-1 and BP10-4-2 failed to produce reads for 10, 10 and 11 genes out of 14, respectively.

**Table 1:** Standard sequencing comparison of library preparation methods by average total number of reads across duplicates, percentage of human and *B. pertussis* reads, average coverage depth of the *B. pertussis* Tohama I reference genome and total coverage of both *B. pertussis* Tohama I and ATCC®9797™ 18323 genomes.

| Library Preparation Method | CFU/mL | Post-depletion *IS481* PCR (C<sub>T</sub>) | Post-depletion *ERV3* PCR (C<sub>T</sub>) | Total Reads | % of Human reads | % of *B. pertussis* reads | Average Coverage Depth | Coverage of the *B. pertussis* Tohama I Genome (%) | Coverage of the *B. pertussis* ATCC Genome (%) |
| --- | --- | --- | --- | --- | --- | --- | --- | --- | --- |
| Nextera XT |  |  |  |  |  |  |  |  |  |
| BP10-2 | 1.4 x 10<sup>6</sup> | 23.53 | 36.48 | 1,465,653 | 40.9% | 0.44% | 1.36 | 57.02% | 56.11% |
| BP10-3 | 1.4 x 10<sup>5</sup> | 28.72 | 36.35 | 3,446,052 | 36.7% | 1.29% | 0.48 | 3.77% | 3.77% |
| BP10-4 | 1.4 x 10<sup>4</sup> | 32.75 | 36.09 | 1,690,120 | 55.8% | 0.30% | 0.18 | 0.77% | 0.78% |
| Illumina DNA prep |  |  |  |  |  |  |  |  |  |
| BP10-2 | 1.4 x 10<sup>6</sup> | Same as above |  | 5,207,437 | 51.0% | 4.25% | 4.25 | 89.16% | 93.28% |
| BP10-3 | 1.4 x 10<sup>5</sup> | Same as above |  | 7,635,405 | 36.4% | 0.89% | 0.89 | 13.31% | 13.78% |
| BP10-4 | 1.4 x 10<sup>4</sup> | Same as above |  | 3,095,938 | 57.8% | 0.37% | 0.01 | 1.26% | 1.27% |
| TruSeq |  |  |  |  |  |  |  |  |  |
| BP10-2 | 1.4 x 10<sup>6</sup> | Same as above |  | Not Sequenced |  |  |  |  |  |
| BP10-3 | 1.4 x 10<sup>5</sup> | Same as above |  | Not Sequenced |  |  |  |  |  |
| BP10-4 | 1.4 x 10<sup>4</sup> | Same as above |  | 514,032 | 67.5% | 0.33% | 0.34 | 0.33% | 0.34% |

### Illumina DNA Library Preparation using standard sequencing

Similar to Nextera XT, the percentage of *B. pertussis* reads recovered decreased with decreasing CFU. However, unlike Nextera XT, genome mapping of BP10-2 showed Illumina DNA prep reads covered 91.2 ± 2.1% of both Tohama I and ATCC®9797™ 18323 genomes, thus, increasing whole genome coverage by 34.7%. Only one sample of BP10-4 was sequenced due to BP10-4-2 failing to pass library QC checkpoints with low library concentrations at the correct size of 300-600 bp. Compared to Nextera XT, Illumina DNA prep resulted in a more even distribution of reads across the vaccine antigens and 23S ribosomal RNA (Figure 1 and Figure 3). BP10-2 Illumina DNA prep at a standard sequencing level was able to provide 4.1 X coverage across all 14 vaccine antigen genes. BP10-3 and BP10-4 produced zero coverage for 6 and 10 genes respectively out of the 14. To test whether the library preparation result could be enhanced by deep sequencing, we sequenced these libraries (BP10-2, BP10-3 and BP10-4) at a concentration estimated to generate 80 million reads.

**Figure 1.**
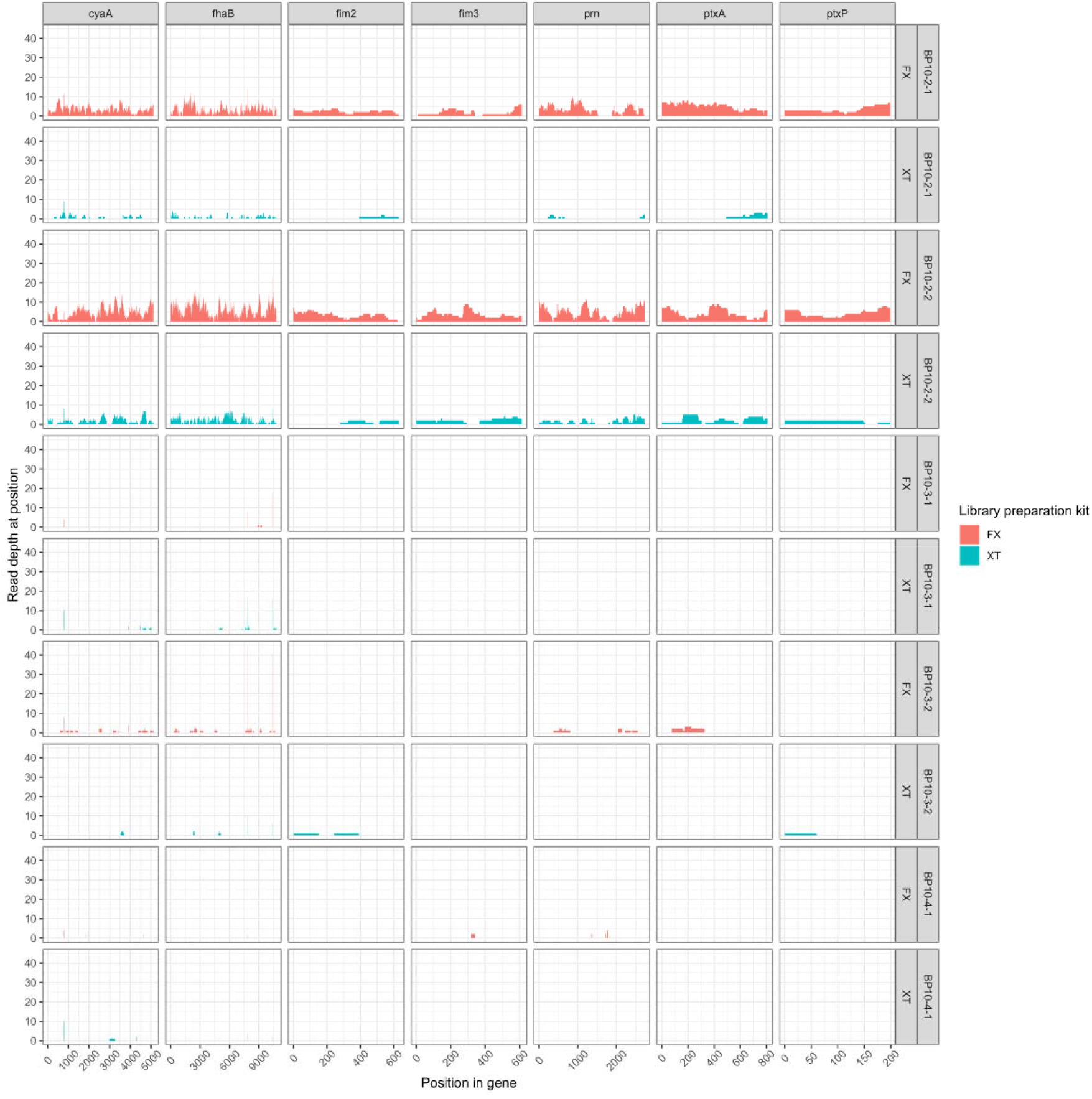
Comparison of Standard Sequencing (SS) gene coverage map over the sequences of the vaccine antigens – *cyaA, fhaB, fim2, fim3, prn, ptxA and ptxP*. The figure also compares the Illumina DNA prep (FX) (Red) and Nextera XT (XT) (Teal) kits at a SS level. The TruSeq (TN) kit was not presented in this figure as no reads were obtained across these regions. Coverage was capped at a maximum of 40x coverage.

### TruSeq DNA Nano Library Preparation using standard sequencing

Most samples of the TruSeq library preparation showed promising library concentration (236 – 3,350 pg/μL), however the majority of the library fragments were only 92-106 bp in size, which were too small to be sequenced. Only BP10-4-2 had a library peak at 490 bp with a concentration of 37.6 pg/μL, hence only this sample was sequenced. BP10-4-2 produced 127,330 reads with 426 reads to *B. pertussis*, these reads provided a depth of 0.34 X across 0.33% of the genome. TruSeq at standard sequencing level of BP10-4-2 failed to yield any reads to genes of interest (Supplementary Material)

### Comparisons of Illumina DNA prep at Standard and Deep Sequencing Levels

Based on the results above, all Illumina DNA prep libraries were re-sequenced on a High-Throughput Flow cell on the NextSeq 500 platform, delivering a total of 493,937,514 reads to 5 samples – BP10-2 (n=2), BP10-3 (n=2) and BP10-4 (n=1). Average read depth and coverage across the genome are presented in Table **2**, and library quantification C_T_ cycles are presented in the Supplementary Material. The number of reads in deep sequencing was amplified 16-18 times and accordingly improved average read depth across the genome by 5-16-fold. In the Illumina DNA prep deep sequencing level, BP10-2 reads provided full 100% coverage of all 14 genes with 50-90X depth, however BP10-3 and BP10-4 had partial coverage only.

**Table 2:** Summary of Illumina DNA prep deep sequencing average reads between duplicate samples.

|  | CFU/m<br>L | Post-depletion<br>*IS481*<br>PCR (C<sub>T</sub>) | Post-depletion<br>*ERV3*<br>PCR (C<sub>T</sub>) | KAPA<br>PCR (C<sub>T</sub>) | Total Reads | % of<br>human<br>DNA<br>reads | % of *B.*<br>*pertussis*<br>DNA reads | Average<br>read<br>depth | Coverage<br>across *B.*<br>*pertussis*<br>Tohama I<br>(%) |
| --- | --- | --- | --- | --- | --- | --- | --- | --- | --- |
| BP10-2 | 1.4 x 10<sup>6</sup> | 23.53 | 36.48 | 15.44 | 82,375,087 | 53.44% | 2.71% | 67.93 | 95.22 |
| BP10-3 | 1.4 x 10<sup>5</sup> | 28.72 | 36.35 | 14.32 | 138,165,091 | 49.76% | 2.05% | 16.07 | 4.68 |
| BP10-4 | 1.4 x 10<sup>4</sup> | 32.73 | 36.09 | 16.32 | 52,857,158 | 60.18% | 0.45% | 3.86 | 1.81 |
| BP12 | N/A | 25.71 | - | 19.04 | 48,289,160 | 27.84% | 2.20% | 20.00 | 98.82 |
| BP16 | N/A | 22.30 | 38.56 | 15.96 | 6,773,770 | 31.18% | 4.14% | 9.70 | 95.68 |

Comparisons of standard and deep sequencing across the vaccine antigen genes of all Illumina DNA prep libraries are presented in Figure 2. Given that a mutation (A **→** G) present in position 2037 of *B. pertussis* Tohama I has been accepted as indicative of macrolide resistance in *B. pertussis*, coverage of this region was investigated (Figure 3). All kits provided coverage over position 2037 ranging from 66 X coverage in BP10-2-1 standard level Nextera XT to 2795 X coverage in BP10-2-1 deep sequencing Illumina DNA prep.

**Figure 2.**
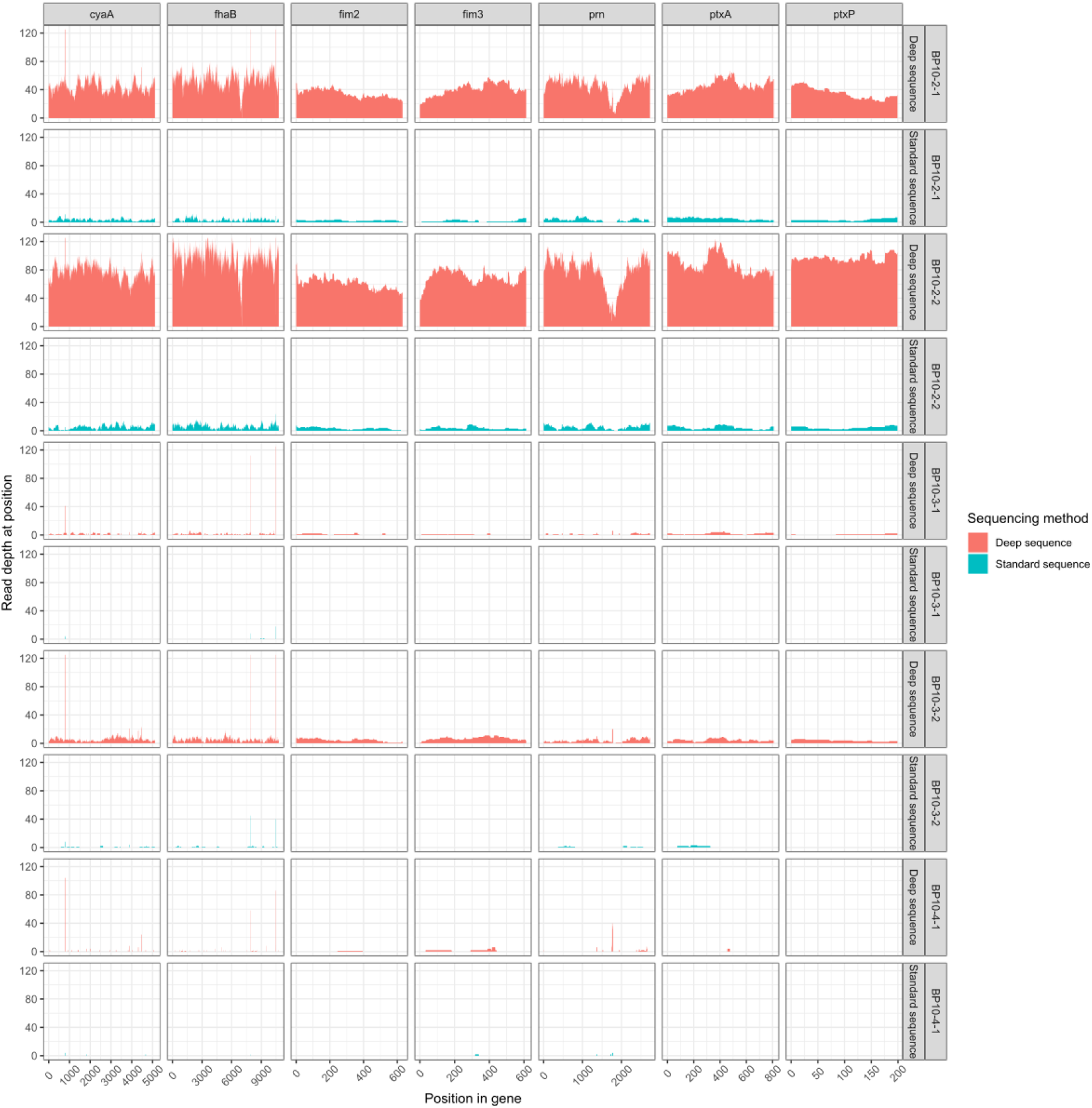
Gene coverage map for genes coding for vaccine antigens. Sequencing at Standard Sequencing (Blue) and Deep Sequencing (Red) levels of Illumina DNA prep (FX) library are shown. Coverage in this figure was capped at 125, only small regions of the genes contained reads exceeding 400X coverage.

**Figure 3.**
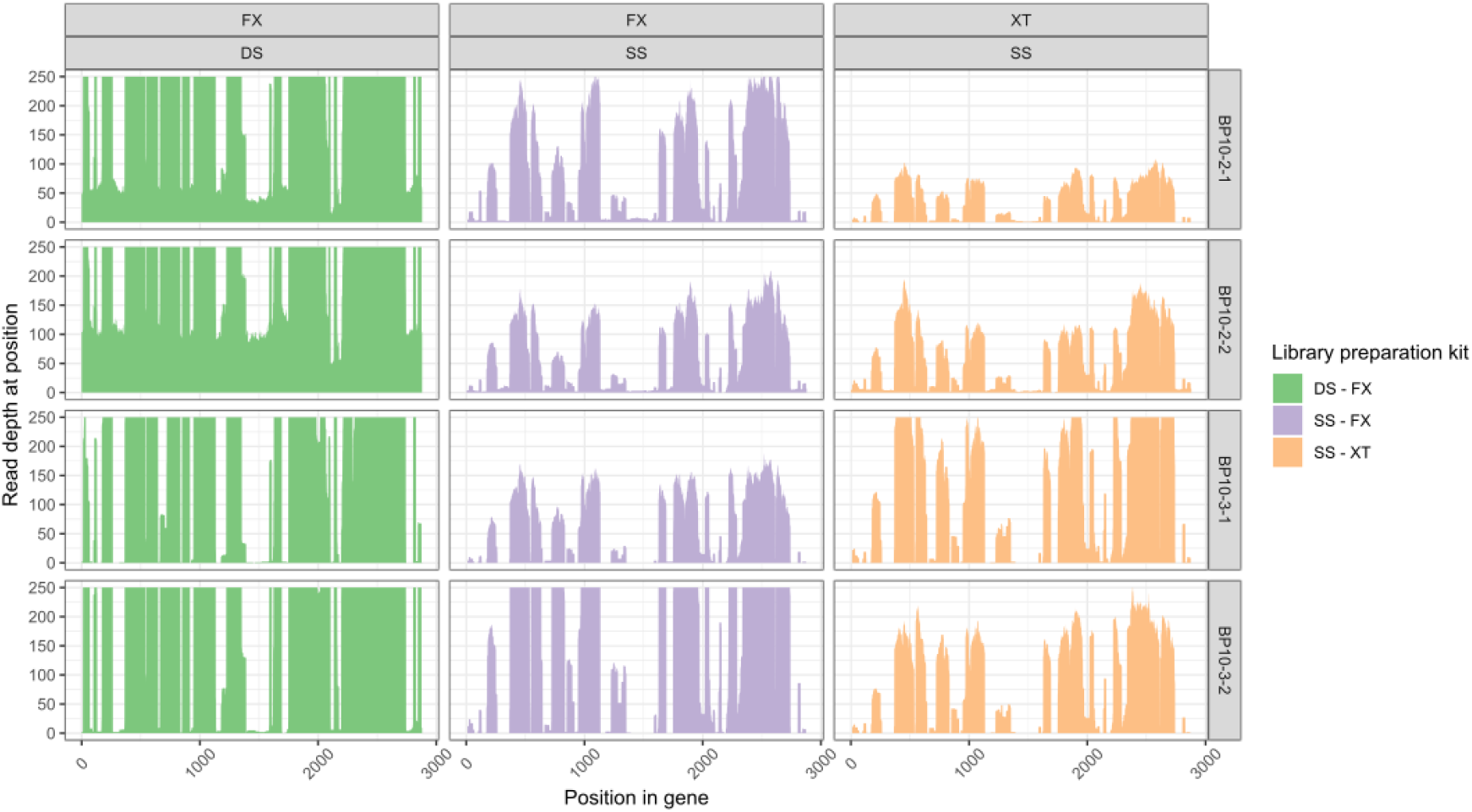
Coverage of the 23S rRNA gene of *B. pertussis*, comparing library preparation kits - Illumina DNA prep (FX) and Nextera XT (XT), and Deep Sequencing (DS) and Standard Sequencing (SS) depths, with varying *B. pertussis* spiked CFU concentrations (BP10-2 1.4 × 10^6^ CFU/mL and BP10-3 1.4 × 10^5^ CFU/mL duplicates).

### Trials on Clinical NPA specimens

Standard sequencing using Illumina DNA prep of the two clinical samples BP12 and BP16, yielded 3,443,046 and 514,032 total reads, with 70,960 (2.06%) and 24,141 (4.69%) *B. pertussis* reads, respectively. These reads produced an average coverage depth of 2.54 X for BP12 and 1.57 X for BP16. Furthermore, BP12 reads covered 50.2% of the B1917 (NZ_CP009751.1) genome, while BP16 covered 42.5%. When these libraries were sequenced at a deeper level the genome coverage increased by +48.7% and +53.2% for BP12 and BP16 respectively Table **2** and Figure 4. Analysis of the genes of interest showed variability in recovery with most genes having more than 92% coverage, with an average depth of 32.91 X and 10.98 X respectively (Supplementary Material).

**Figure 4.**
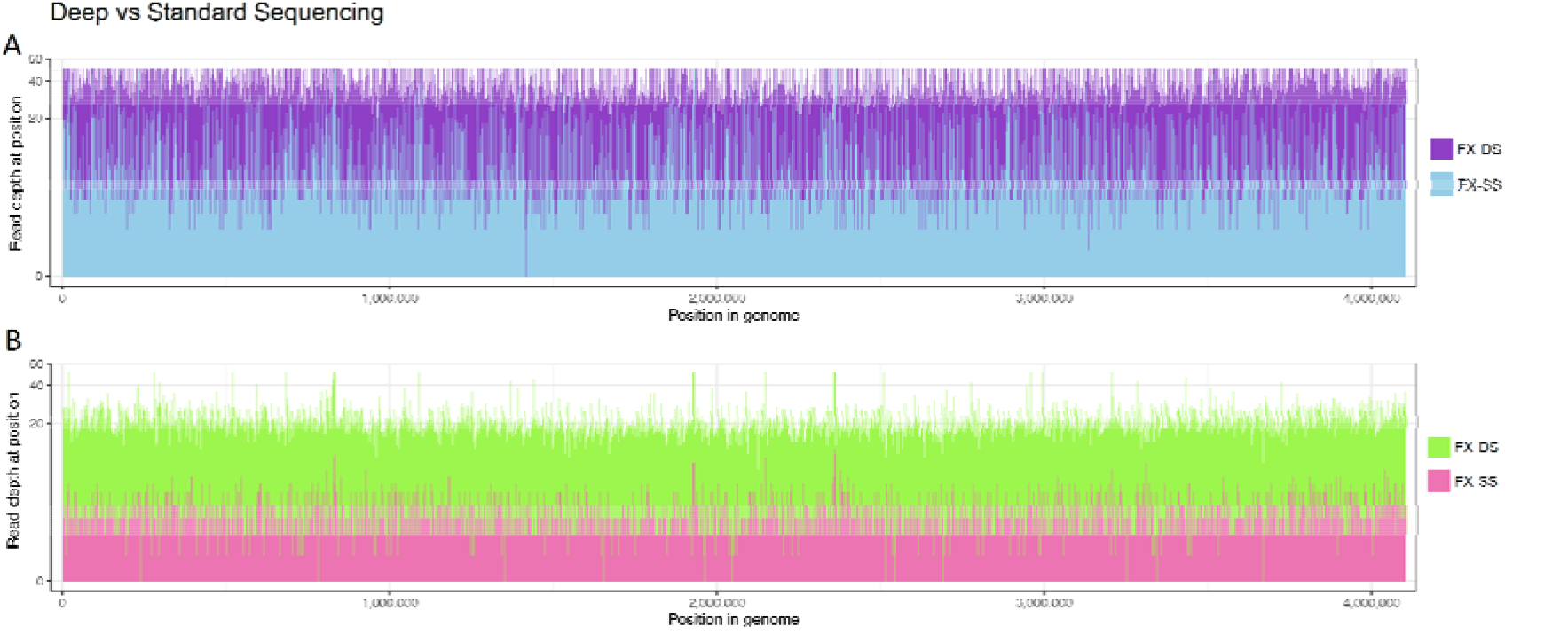
Whole genome coverage map of the positive strand in BP12 and BP16 at Standard Sequencing (SS) (Blue and Pink) and Deep Sequencing (DS) Levels (Purple and Green), mapped to B1917 (NZ_CP009751.1). Panel A represents BP12 and Panel B represents BP16. Scale of both samples have been capped at 50 X depth.

## Discussion

In the era of culture-independent diagnostic testing, the primary consequence is the loss of routine laboratory culture. Without cultured isolates, many gold-standard phenotypic typing systems have been lost. Therefore, further laboratory classification of currently circulating strains has become unavailable and this in turn affects public health outbreak control strategies. The aim of this study was to compare commercially available library preparation methods and sequencing depths to enhance our ability to directly sequence *Bordetella pertussis-*positive nasopharyngeal aspirates. Comparisons were made based on sequencing read depths, average genome coverage and the ability to accurately extract sequencing data from regions of interest such as the vaccine antigens.

This study demonstrated that different library preparation methods can impact the recovery of important genomic information from culture-independent sequencing. Nextera XT libraries at a standard sequencing depth produced uneven coverage across the genome, only accounting for ∼50% of the *B. pertussis* Tohama I genome, compared to more than 80% of the genome when libraries are prepared with Illumina DNA prep. Overall, the Illumina DNA prep library preparation kit generated better average coverage across the *B. pertussis* genome at both the standard and deep sequencing depths. Sequencing at a deeper level only amplified depth and increased confidence to detect SNPs. In line with what others have shown^4^, we found that sequencing at a standard level with any library preparation method, was insufficient in obtaining accurate molecular typing information.

We report variations in total reads between all Illumina DNA prep samples most likely due to library normalisation and final library pooling. Interestingly, we noted that in this study the library concentrations ranged between 3 C_T_ cycles from each other and these C_T_ cycles reflected how many sequencing reads were dedicated to each sample. This variation is generally acceptable at a standard sequencing scale, hence further investigations into the effect of library concentration and normalisation would need to be performed to optimise deep sequencing protocols and ensure an even distribution of sequencing resources for all samples.

The TruSeq DNA Nano library kit was chosen based on its fragmentation method – mechanical instead of enzymatic to reduce the inherent G+C bias of the transposase. However, post-library preparation QC determined very poor concentrations of library fragments in the appropriate range (400-500bp). TruSeq libraries potentially failed due to low starting concentration of DNA given the sample had been host DNA depleted. Fragment size analysis was performed before and after sonication on BP10-2 samples, however low input DNA concentration made it difficult to observe fragment size peaks efficiently. Sequencing of one of the TruSeq libraries (BP10-4-2) proceeded due to the presence of a peak in the 400-500bp region. TruSeq libraries were constructed with other samples with higher starting DNA concentration, which sequenced successfully, hence library preparation methods and reagents were not the problem. Therefore, TruSeq was not an appropriate kit for low DNA concentrations as selection of sonication parameters is difficult to optimise for host depleted clinical samples with low concentration of DNA.

As expected, decreasing *B. pertussis* CFU in spiked samples were able to demonstrate that changes in bacterial load can influence the amount of sequencing reads recovered from a sample. Based on routine rtPCR results, clinical *IS481-*positive specimens have an average C_T_ cycle of 30.92, suggesting a CFU load of 1.4 × 10^4^ (CFU/mL) equivalent to sample BP10-4, is a typical *B. pertussis* load for clinical NPA samples. As such, the higher number of *B. pertussis* cells in BP10-2 of 1.4 ×10^6^ CFU is an uncommon occurrence in a clinical sample. The two clinical cases enrolled in this study had a very high initial C_T_ cycle of 12.15 (BP12) and 17.97 (BP16), and their total read counts of 48,289,160 and 6,773,770, respectively, demonstrate the importance of quality clinical sample collection ensuring high load of target bacterial DNA prior to sequencing to allow sufficient genome coverage.

To achieve adequate depth and coverage of the *B. pertussis* genome in a low bacterial load NPA sample would require further enrichment of the clinical sample and/or library. However, our results indicate that a *B. pertussis* positive sample with a CFU equivalent to or more than 1.4 ×10^6^ CFU would, without enrichment, generate coverage and depth sufficient for genomic surveillance analyses. By combining host DNA depletion protocols (Saponin), an appropriate library preparation method (Illumina DNA prep) and applying a deep sequencing approach (>50 million reads) with a library quantification result of more than 15 C_T_ cycles, should result in 20X coverage at least, across the regions of interest if not the whole genome of *B. pertussis*. While sequencing at a deeper level costs more compared to sequencing a pure isolate, it is now an option when there are few isolates to sequence in this era of PCR diagnostics.

This study was limited by the number of replicates that could be performed as a result of cost-limitations; hence results are subjected to sample bias. Numbers of NPA samples are also slowly declining as preferences to nasopharyngeal swabs are more popular. The volume of liquid in NPA are also very limited due it its use in previous diagnostic protocols and therefore replicates of clinical samples could not be performed. Trialling the protocol on more common respiratory sample types such as nasopharyngeal and throat swabs would also expand its use.

Future research should focus on improving sequencing depth and coverage over regions of interest, particularly genes coding for vaccine antigens and other molecular typing targets. Despite the expense, further deep sequencing trials on larger number of clinical NPA are required to validate the protocol and determine information yield across sample types.

In conclusion, our study demonstrated the feasibility of direct sequencing of *B. pertussis* from clinical NPA specimens with a high bacterial load and the recognition of potentially actionable targets in *B. pertussis* genome. This can be achieved through the combination of optimised sample and library preparation followed by deep sequencing.

## Author Statements

The study was conceptualised by WF, VT, and VS. Laboratory work was performed by WF, RR and VT. Bioinformatics and genome analysis was executed by WF, KP and RS. The manuscript was written by WF and reviewed and edited by KP, RR, RS, VT and VS.

The authors declare no conflict of interest

Reads mapping to *Bordetella pertussis* ATCC9797 18323 for spiked nasopharyngeal aspirates and BP1917 for clinical nasopharyngeal aspirates have been uploaded to SRA under BioProject: PRJNA694997

This study as supported financially by the Centre for Infectious Diseases and Microbiology – Public Health Post-graduate scholarship

Consent for the images in this article has been provided by the authors

This work was supported by the Prevention Research Support Program, funded by the New South Wales Ministry of Health. Special thanks to Illumina for providing a complimentary Illumina DNA prep Library Preparation Kit. The authors are grateful to the staff of Centre for Infectious Diseases and Microbiology Laboratory Services, NSW Health Pathology for their technical assistance and expertise. Computational analysis was performed on the University of Sydney High Performance Cluster, with the assistance of the Sydney Informatics Hub. The authors would like to acknowledge the Microbial Genomics Reference Laboratory, Centre for Infectious Disease and Microbiology –Public Health, Westmead Hospital, for their assistance with genome sequencing and bioinformatics analysis.

Nasopharyngeal aspirates were collected by the Centre for Infectious Diseases and Microbiology Laboratory services under the Western Sydney Local Health District Research Ethics and Governance committee. Project identifier: 2019/PID02294

Consent was not obtained from patients, as these NPA were left-over from previous diagnostic testing, and otherwise discarded. These NPA samples were pooled to prevent any identification during sequencing, and no record of identifiable data was collected.

